# Correction of eIF2-dependent defects in brain protein synthesis, synaptic plasticity and memory in mouse models of Alzheimer’s disease

**DOI:** 10.1101/2020.10.19.344564

**Authors:** Mauricio M. Oliveira, Mychael V. Lourenco, Francesco Longo, Nicole P. Kasica, Wenzhong Yang, Gonzalo Ureta, Danielle D. P. Ferreira, Paulo H. J. Mendonça, Sebastian Bernales, Tao Ma, Fernanda G. De Felice, Eric Klann, Sergio T. Ferreira

**Affiliations:** Institute of Medical Biochemistry Leopoldo de Meis, Federal University of Rio de Janeiro, Rio de Janeiro, RJ, Brazil; Center for Neural Science, New York University, New York, NY, USA; Department of Internal Medicine, Gerontology & Geriatric Medicine, Wake Forest School of Medicine, Winston-Salem, NC, USA; Fundación Ciencia & Vida, Santiago, Chile; Institute of Biomedical Sciences, Federal University of Rio de Janeiro, Rio de Janeiro, RJ, Brazil; Centre for Neuroscience Studies and Department of Psychiatry, Queen’s University, Kingston, ON, Canada; NYU Neuroscience Institute, New York University School of Medicine, New York, NY, USA; Institute of Biophysics Carlos Chagas Filho, Federal University of Rio de Janeiro, Rio de Janeiro, RJ, Brazil

**Keywords:** Alzheimer’s disease, integrated stress response, ISRIB, protein synthesis, eIF2 alpha, mRNA translation

## Abstract

Neuronal protein synthesis is essential for long-term memory consolidation. Conversely, dysregulation of protein synthesis has been implicated in a number of neurodegenerative disorders, including Alzheimer’s disease (AD). Several types of cellular stress trigger the activation of protein kinases that converge on the phosphorylation of eukaryotic translation initiation factor 2α (eIF2α-P). This leads to attenuation of cap-dependent mRNA translation, a component of the integrated stress response (ISR). We show that AD brains exhibit increased eIF2α-P and reduced eIF2B, key components of the eIF2 translation initiation complex. We further demonstrate that attenuating the ISR with the small molecule compound ISRIB (ISR Inhibitor) rescues hippocampal protein synthesis and corrects impaired synaptic plasticity and memory in mouse models of AD. Our findings suggest that attenuating eIF2α-P-mediated translational inhibition may comprise an effective approach to alleviate cognitive decline in AD.

## Introduction

Alzheimer’s disease (AD) is the most prevalent form of dementia, affecting more than 35 million people worldwide. Pathological hallmarks of AD include brain accumulation of the amyloid-β peptide (Aβ) and of neurofibrillary tangles composed of hyperphosphorylated tau protein, neuroinflammation, synapse failure/loss, and neurodegeneration, ultimately culminating in memory failure (*1*). Despite major progress in the elucidation of mechanisms of pathogenesis in recent years, AD is still a disease in urgent need of effective therapeutics capable of preventing and/or blocking progressive cognitive deterioration. Given the complex nature of AD, identification of molecular pathways and targets that effectively improve cognition has been challenging.

De novo protein synthesis plays a key role in synaptic plasticity and long-term memory consolidation (*2–7*). Initiation of cap-dependent mRNA translation involves the formation of a ternary complex comprising Met-tRNA, GTP and the eukaryotic initiation factor 2 (eIF2) complex. Activity of eIF2B, a guanine exchange factor (GEF), allows continuous ribosome assembly and mRNA translation (*8*). This step is tightly regulated by phosphorylation of eIF2 on its alpha subunit (eIF2α-P), which inhibits the GEF activity of eIF2B and attenuates global translation (*9, 10*).

Various cellular stress stimuli trigger activation of eIF2α kinases causing phosphorylation of eIF2α in a process termed the Integrated Stress Response (ISR). The ISR attenuates global cap-dependent protein synthesis while favoring the translation of a specific subset of mRNAs that help to restore cellular homeostasis (*11*). Increased translation of activating transcription factor 4 (ATF4, also known as CREB-2) is thought to be a central mediator of the ISR (*12*).

Mounting evidence implicates aberrant ISR and eIF2α-P in brain disorders, including traumatic brain injury, prion disease, Down syndrome, and AD (*13–23*). ISR markers are elevated in AD brains (*16, 24, 25*), as well as in the brains of mouse models of AD (*15, 16, 26*). We previously demonstrated that either genetic ablation or inhibition of eIF2α kinases rescues synapse function and memory defects in AD mouse models (*15, 16*). We thus hypothesized that rescuing brain protein synthesis downstream of eIF2α-P might be an attractive approach to correct cognitive impairment in AD.

Herein, we report that AD brains exhibit increased eIF2α-P levels accompanied by reductions in eIF2B subunits. The ISR inhibitor, ISRIB, a small molecule compound that stimulates eIF2B activity even in the presence of elevated eIF2α-P (*27–29*), rescued reduced hippocampal protein synthesis and corrected impaired synaptic plasticity and memory in AD model mice. Our findings suggest that targeting dysregulated brain translational control by eIF2α may represent an effective approach to combat cognitive failure in AD.

## Results

### AD brains present increased eIF2α-P and reduced eIF2B subunit levels

We first confirmed that cortical extracts from AD brains (see Supplementary Table 1 for demographics) displayed increased eIF2α-P (Fig. 1A), consistent with previous findings (*16, 24*). No changes in total levels of the α and γ subunits of eIF2 were detected (Fig. 1B,C). In contrast, both α (required for assembly) and ε (catalytic) subunits of eIF2B were substantially reduced in AD (Fig. 1D,E). These results indicate that altered translation initiation in AD brains is associated with ISR activation (as assessed by eIF2α-P) and with reductions in eIF2B subunits. Because mRNA translation is central to long-term memory consolidation, these findings suggest that rescuing brain eIF2-mediated translation could be a target to treat memory failure in AD.

**Figure 1.**
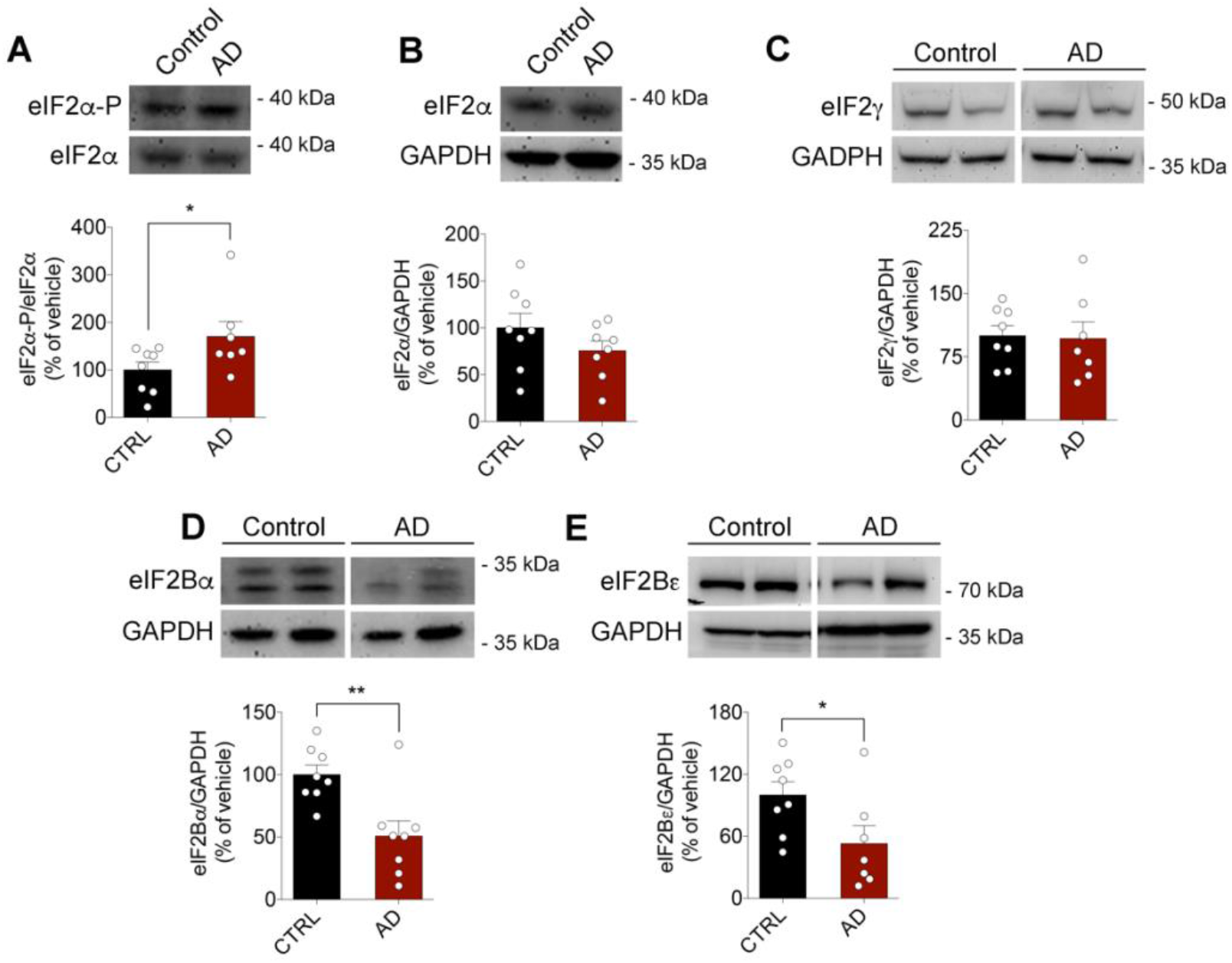
AD brains display increased eIF2α-P and reduced levels of eIF2B subunits. Cortical tissue from AD subjects and age-matched controls were probed for translation initiation factors. Samples were probed for eIF2α-P (A; F = 3.125; N = 7 Control cases; N = 7 AD cases), total eIF2α (B; F = 1.319; N = 8 Control cases; N = 8 AD cases), eIF2γ (C; F = 1.547; N = 8 Control cases; N = 7 AD cases), eIF2Bα (D; F = 1.375; N = 8 Control cases; N = 8 AD cases) and eIF2Bε (E; F = 2.823; N = 8 Control cases; N = 7 AD cases) by Western blotting (AD). eIF2α-P was normalized by total eIF2α. eIF2α, eIF2γ, eIF2Bα and eIF2Bε were normalized by GAPDH. Non-contiguous lanes from the same blot are shown in C, D and E. Outliers were excluded from each analysis using Grubb’s test (α = 0.05) from GraphPad Prism 6. *p < 0.05; **p < 0.01; Two-tailed unpaired *t*-test. Dots represent individual subjects.

### ISRIB prevents eIF2α-P-induced impairment of long-term memory in mice

We performed a proof-of-concept study to determine whether the ISR inhibitor, ISRIB, could prevent memory impairment caused by brain accumulation of eIF2α-P. ISRIB has been shown to bind to and stabilize the eIF2B complex in its active form, thereby bypassing eIF2α-P to stimulate the free eIF2B pool (*29*). We first determined whether systemic (i.p.) administration of ISRIB resulted in its accumulation in the mouse brain. To this end, we administered ISRIB (0.25 mg/kg, i.p.) to mice for six consecutive days and measured ISRIB concentrations in plasma and brain extracts by mass spectrometry. Results showed that ISRIB could be detected in both plasma and brain at 4- and 24-hours following drug administration (Fig. 2A).

**Figure 2.**
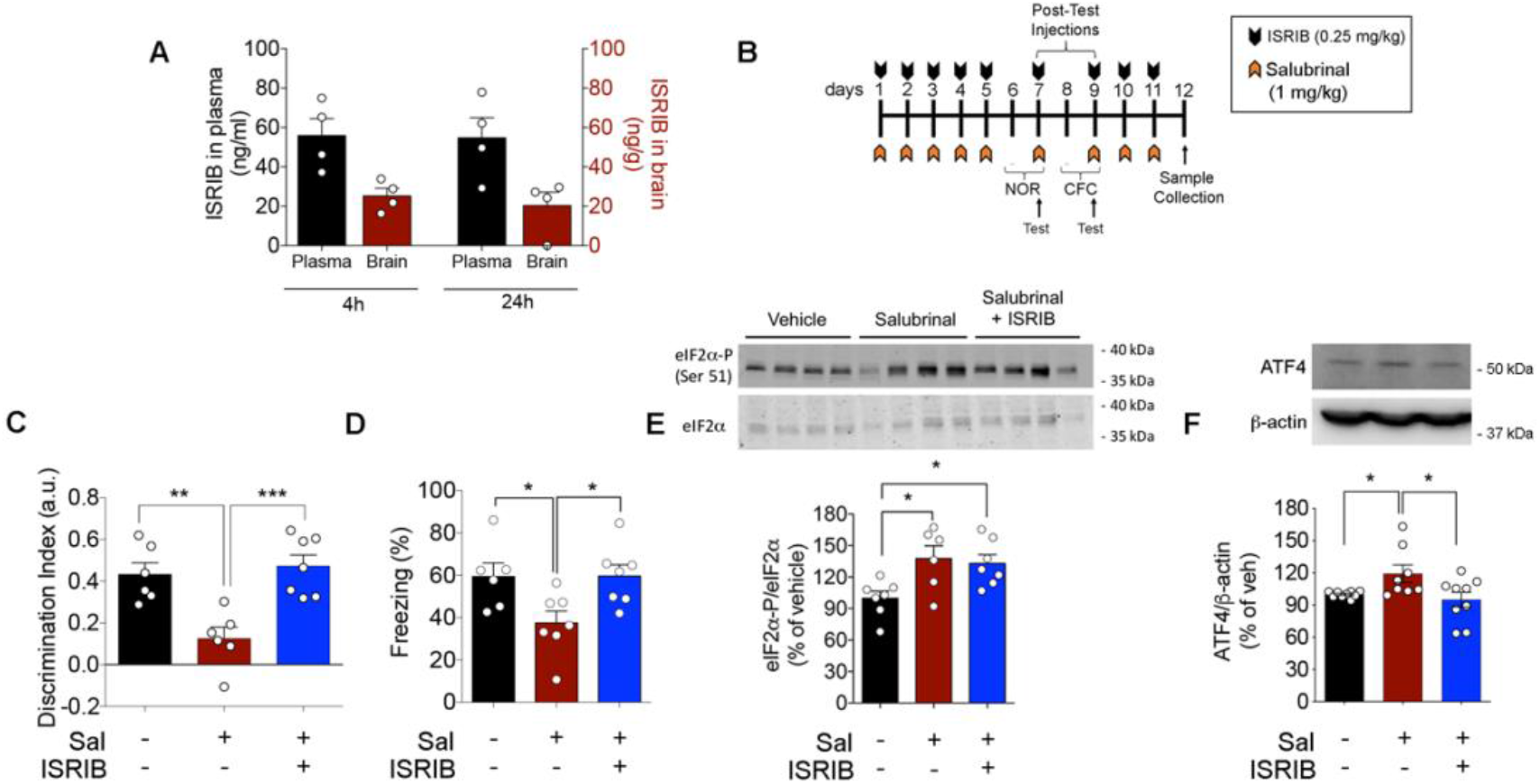
Systemic treatment with ISRIB prevents memory impairment induced by eIF2α-P. (A) ISRIB concentrations in plasma and brain were determined by mass spectrometry 4 or 24 hours after i.p. administration (0.25 mg/kg for 6 consecutive days) in mice (N = 4 mice/time point). (B) Experimental timeline: C57BL/6 mice were treated with salubrinal alone (10 mg/kg, i.p.; orange arrows) or salubrinal + ISRIB (0.25 mg/kg, i.p.; black arrows) on days 1-5, 7, 9-11. Mice were trained and tested in the Novel Object Recognition (NOR) task on days 6 and 7, respectively, and were trained and tested in the Contextual Fear Conditioning (CFC) task on days 8 and 9, respectively. Brains were collected on day 12. (C) Discrimination index in the NOR test (F = 12.00; N = 5-7 mice/group;). (D) Freezing time in the CFC test (F = 5.049; N = 5-7 mice/group). (E) Hippocampal eIF2α-P in mice treated with salubrinal alone or salubrinal + ISRIB, normalized by total eIF2α (N = 6-7 mice/group; F = 5.461). In all panels, dots correspond to individual mice. *p < 0.05, **p < 0.01, ***p<0.001, One-way ANOVA followed by Dunnet’s *post hoc* test.

In another set of experiments, C57BL/6 mice were given daily injections of salubrinal (1 mg/kg, i.p., for 5 consecutive days; Fig. 2B), an inhibitor of eIF2α dephosphorylation). Long-term memory was then assessed using the Novel Object Recognition (NOR) and Contextual Fear Conditioning (CFC) tasks. Treatment with salubrinal impaired long-term memory in both NOR (Fig. 2C) and CFC (Fig. 2D) tests. Systemic treatment of mice with ISRIB (0.25 mg/kg, i.p.) prevented memory impairments induced by salubrinal in both NOR and CFC tests (Figs. 2C-D). ISRIB did not prevent the increase in hippocampal eIF2α-P caused by salubrinal administration in mice (Fig. 2E) but blocked the accumulation of ATF4, a central mediator of the ISR (Fig. 2F). These findings are consistent with the notion that ISRIB acts downstream of eIF2α-P to restore memory, and indicate that systemically administered ISRIB reaches the brain and prevents memory impairments induced by the accumulation of eIF2α-P.

### ISRIB prevents eIF2α-P-mediated impairment in long-term memory in an acute model of Alzheimer’s disease

We next asked whether ISRIB could counteract the activation of ISR and memory deficits that are induced by amyloid-β oligomers (AβOs), toxins that accumulate in AD brains and cause eIF2α-P-mediated synapse and memory failure (*1, 15, 30*). To investigate this possibility, C57BL/6 mice received a single intracerebroventricular (i.c.v.) infusion of AβOs (10 pmol, expressed as Aβ monomers; (*15, 31, 32*) followed by administration of ISRIB (0.25 mg/kg, daily, i.p.) for the duration of the experiment (Fig. 3A). Because ISRIB has been reported to trigger pro-mnemonic effects (Sidrauski et al., 2013), we did not inject ISRIB between training and test sessions of memory tasks. I.c.v. infusion of AβOs induced hippocampal eIF2α-P (Fig. 3B) and increased ATF4 levels (Fig. 3C). Treatment with ISRIB counteracted the increase in hippocampal ATF4 protein levels (but not mRNA; Figs. 3C and S1A) without affecting eIF2α-P levels (Fig. 3B).

**Figure 3.**
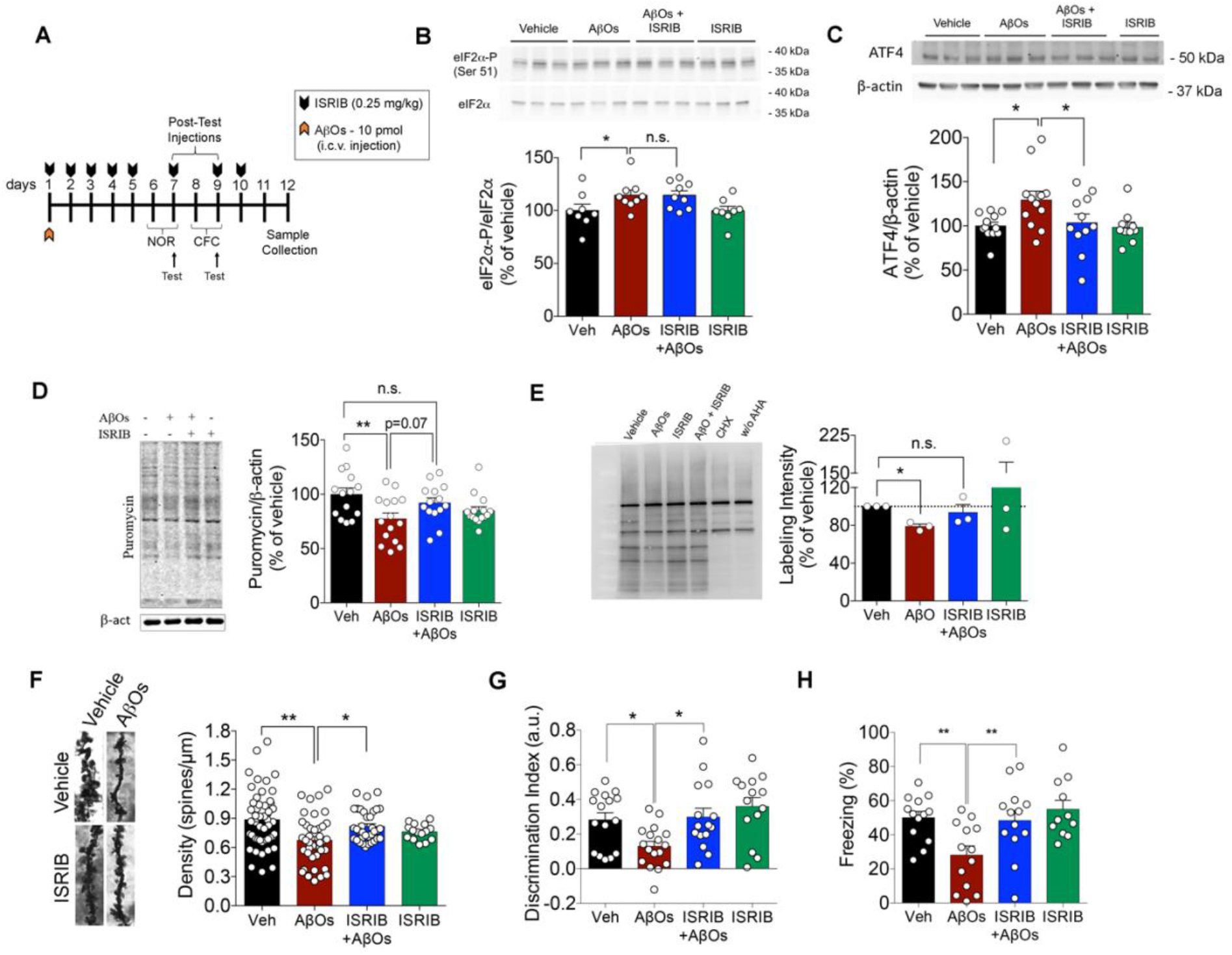
ISRIB prevents memory impairments, dendritic spine loss and defective hippocampal protein synthesis induced by AβOs. (A) Experimental timeline: C57BL/6 mice received an i.c.v. infusion of AβOs (10 pmol; orange arrow) and were treated with ISRIB (0.25 mg/kg, i.p.; black arrows) or saline on days 1-5, 7, 9-11. Mice were trained and tested in the Novel Object Recognition (NOR) task on days 6 and 7, respectively, and were trained and tested in the Contextual Fear Conditioning (CFC) task on days 8 and 9, respectively. Brains were collected on day 12. (B) eIF2α-P in the hippocampi of mice that received an i.c.v infusion of AβOs (or vehicle) and were treated with ISRIB (0.25 mg/kg, i.p., for 5 days) or saline (N = 8-9 mice/group). (C) ATF4 in the hippocampi of mice that received an i.c.v infusion of AβOs (or vehicle) and were treated with ISRIB (0.25 mg/kg, i.p., for 5 days) or saline (N = 10-12 mice/group). (D) Mouse hippocampal slices were exposed to AβOs (1 μM) in the absence or presence of ISRIB (0.2 μM) for 3h and protein synthesis was measured using SUnSET (N = 15-18 slices from a total of 15 mice per experimental condition). Representative images shown had brightness linearly adjusted for clearer visualization. (E) Primary hippocampal cultures were exposed to AβOs (0.5 μM) in the absence or presence of ISRIB (0.2 μM) for 3h, and newly synthesized proteins were detected by BONCAT (labeling with AHA followed by Click chemistry for biotinylation of AHA-containing polypeptides; see “Methods”). After pulldown with streptavidin-conjugated resin, proteins were detected by Western blotting. Symbols represent experiments with independent hippocampal cultures and independent AβO preparations (N = 3 independent primary cultures). (F) Dendritic spine density was analyzed in apical dendrites of neurons from the CA1 region of the hippocampus. Each symbol corresponds to the mean of three independent 20 μm segments per neuron, 5 neurons per mouse (N = 4-5 mice/group). (G) Discrimination index in the NOR test (F = 5.468; N = 14-17 mice/group). (H) Freezing time in the CFC test (F Interaction = 5.683; N = 11-13 mice/group). *p < 0.05, **p < 0.01, Two-way ANOVA followed by Dunnet’s *post hoc* test for all experiments, except in panels 3I and 3J, analyzed by One-Way ANOVA followed by Dunnet’s *post-hoc* test. Dots represent individual mice.

We then hypothesized that ISRIB could correct translation defects induced by AβOs. To test this hypothesis, we exposed ex vivo mouse hippocampal slices to AβOs (1 μM) in the absence or presence of ISRIB (0.2 μM) and assessed de novo protein synthesis using SUnSET (*33*). We found that ISRIB prevented hippocampal translational repression induced by AβOs (Fig. 3D). Similar results were obtained in primary neuronal cultures using BONCAT (*34*) (Fig. 3E).

We further found that dendritic spine density was reduced in hippocampal CA1 in mice that received an i.c.v. infusion of AβOs, and that systemic treatment with ISRIB restored spine density in AβO-infused mice (Fig. 3F). i.c.v. infusion of AβOs caused reductions in hippocampal synaptophysin, PSD-95 and BDNF, and treatment with ISRIB had no effect on these proteins (Fig. S1B-E).

We next tested whether systemic treatment with ISRIB could prevent AβO-induced cognitive impairment. We found that ISRIB prevented AβO-induced long-term memory failure in both NOR (Fig. 3G) and CFC (Fig. 3H) tasks. Control measurements showed no differences in locomotor or exploratory activities in mice that received an AβO infusion and/or were treated with ISRIB (Fig. S2A-C), and no differences in the training phase in either NOR or CFC tests (Fig. S2D,E). These findings indicate that ISRIB corrects AβO-instigated eIF2α-P-dependent hippocampal translational repression, attenuates dendritic spine loss, and rescues memory impairments in mice.

### ISRIB restores synapse function and memory in a transgenic mouse model of AD

Next, we investigated whether ISR inhibition could reverse deficits in synaptic plasticity and memory in 10-13 month-old APPswe/PS1ΔE9 mice, a transgenic mouse model of AD characterized by age-dependent brain accumulation of Aβ (*35*). We initially investigated the effect of ISRIB on long-term potentiation (LTP) in hippocampal slices from APPswe/PS1ΔE9 mice. Whereas slices from APPswe/PS1ΔE9 mice failed to maintain LTP at CA1 Schäffer collateral synapses following tetanic stimulation, treatment with ISRIB restored LTP (Fig. 4A-B). We further found that ISRIB restored spine density in pyramidal neurons from hippocampal CA1 region of APPswe/PS1ΔE9 (Fig. 4C).

**Figure 4.**
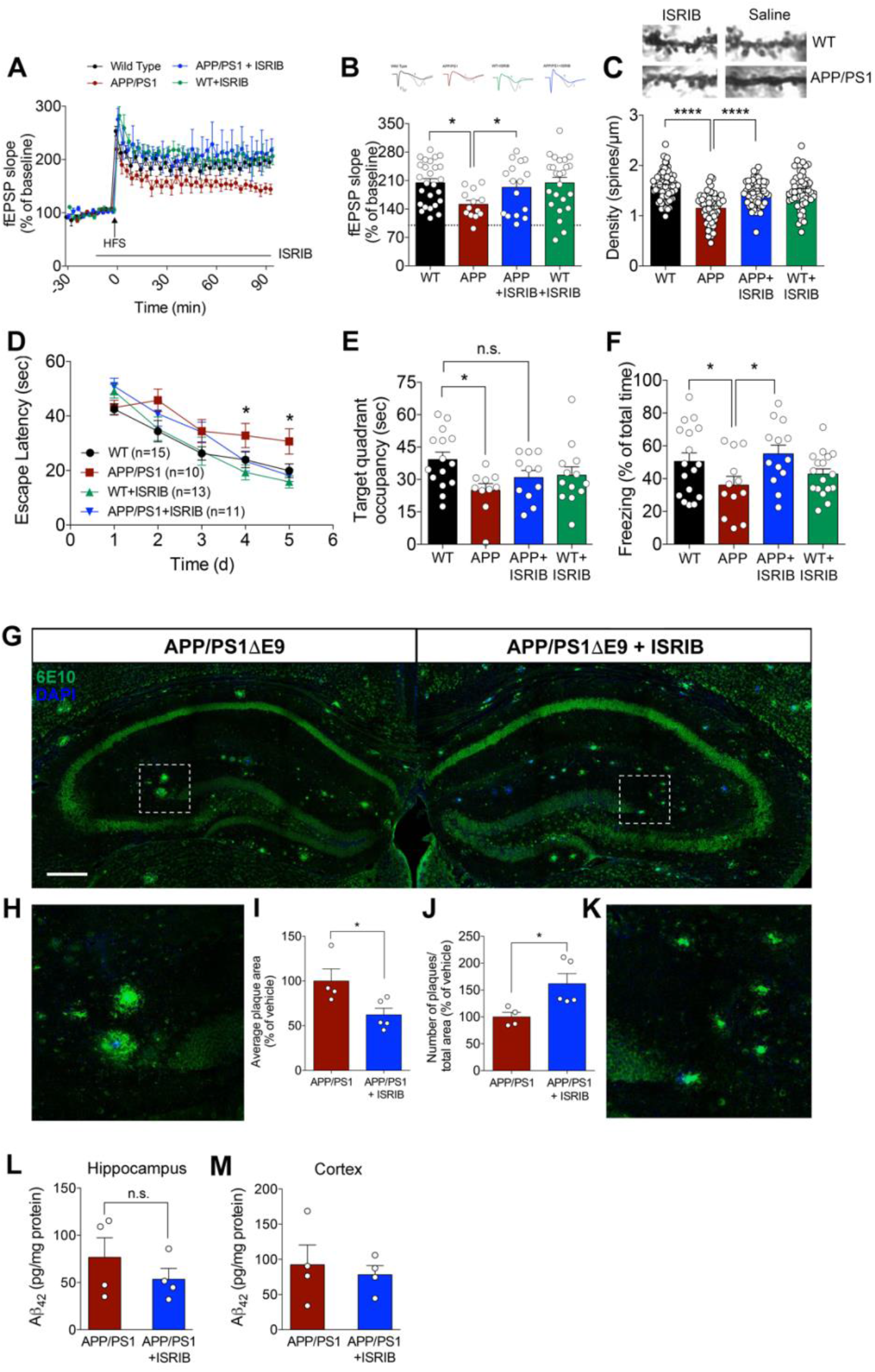
ISRIB reverses defective hippocampal LTP and memory in APPswe/PS1ΔE9 mice. 10-13 month-old APPswe/PS1ΔE9 mice (or WT littermates) were treated daily for 2 weeks with ISRIB (0.25 mg/kg, i.p.) prior to behavioral analysis. (A) Field excitatory post-synaptic potential recordings in acute hippocampal slices from WT or APPswe/PS1ΔE9 mice exposed to vehicle or 0.2 μM ISRIB (N = 13-26 slices from 6-9 mice/group). (B) On top - Representative slopes for fEPSP measurements. “1” is representative of the baseline, and “2” is representative of potentiation. Graph is representative of field excitatory post-synaptic potentials 90 min after tetanic stimulation. (C) Dendritic spine density was analyzed in apical dendrites of neurons from the CA1 region of the hippocampus. Each dot is representative of one segment using in the analysis (N = 4 mice/group) (D) Mice were trained for 5 days in the MWM, and latency time to reach the platform was recorded (N = 10-15 mice/group). Symbols represent means ± SE of the latency times to find the platform on each consecutive day of training. (E) On the 6^th^ day, the platform was removed, and mice were allowed to freely explore the pool for 1 minute. Total time spent in the target quadrant during the probe trial was measured (N = 10-15 mice per group). (F) Contextual fear conditioning of WT or APPswe/PS1ΔE9 mice treated with ISRIB (N = 10-15 mice/group) or saline. (G) Representative photomontage illustrating presence of plaques (6E10 immunoreactivity; green channel) in the dorsal hippocampus of APPSwe/PS1ΔE9 mice treated with vehicle (left) or with ISRIB (right) (N = 4-5 mice/group; scale bar = 200 μm). (H, K) Optical zoom images of regions identified by dashed rectangles in panel G. (I) Average plaque area in the hippocampi of APPswe/PS1ΔE9 mice treated with vehicle or ISRIB (N = 4-5 mice/group). (J) Total number of plaques normalized by hippocampal area. (L, M) Total Aβ_42_ measured by ELISA in hippocampal (L) or cortical (M) homogenates of APPSwe/PS1ΔE9 mice treated with vehicle or ISRIB (N = 4 mice/group). *p < 0.05, ****p < 0.0001 Two-way ANOVA followed by Dunnet’s *post-hoc* test, except for panel 1A, evaluated by Repeated Measures Two-Way ANOVA, followed by Tukey’s *post hoc* analysis. Dots represent individual mice.

To determine the effect of ISRIB on long-term spatial memory, we treated APPswe/PS1ΔE9 mice (or WT littermates) with ISRIB (0.25 mg/kg, i.p.) daily for two weeks before testing in the Morris water maze (MWM). This protocol was found to alleviate the ISR in the hippocampus, as assessed by total levels of GADD34 (Fig. S3). ISRIB improved learning in APPswe/PS1ΔE9 mice, as indicated by faster learning curves compared to saline-treated APPswe/PS1ΔE9 mice (Fig. 4D), but did not improve memory retention assessed in the MWM probe trial (Fig. 4E). However, ISRIB improved long-term contextual memory of APPswe/PS1ΔE9 mice in the CFC test (Fig. 4F). Control experiments showed that APPswe/PS1ΔE9 (or WT) mice treated with ISRIB had no changes in locomotor or exploratory activities, and their weights were similar to control animals (Fig. S4A-D). Results thus showed that treatment with ISRIB restores synaptic plasticity and memory in APPswe/PS1ΔE9 mice.

We proceeded to determine whether treatment with ISRIB affected amyloid deposition in the brains of APPswe/PS1ΔE9 mice. We found that ISRIB-treated APPswe/PS1ΔE9 mice had reduced mean amyloid plaque size (Fig. 4G-I), but increased plaque density in the hippocampal formation (Fig. 4J-K). Total Aβ_42_ in the hippocampus (Fig. 4L) and cortex (Fig. 4M) was unchanged by treatment. We hypothesized that the reduced mean plaque size in APPswe/PS1ΔE9 mice treated with ISRIB could be due to altered plaque phagocytosis by glial cells. APPswePS1ΔE9 mice exhibited increased hippocampal immunoreactivities for Iba-1 (microglial marker) and GFAP (astrocytic marker) compared to WT mice (Fig. S5). Treatment with ISRIB had no effect on either Iba-1 (Fig. S5A-C) or GFAP (Fig. S5D-F) immunoreactivities in the hippocampi of APPswe/PS1ΔE9 mice. Collectively, our findings indicate that ISRIB attenuates translational repression, restores synaptic plasticity and memory independently of amyloid burden in AD models.

## Discussion

Activation of the ISR and attenuation of brain protein synthesis have been implicated in memory deficits in AD (*15, 16, 24–26*) and in other neurodegenerative disease (*13–23*), supporting the notion that correcting defective brain protein synthesis might comprise an effective therapeutic target in AD. Because three out of four known eIF2α kinases have been shown to play pathogenic roles in AD (*15, 16, 24, 25, 36, 37*), identifying approaches that act downstream of eIF2α-P may comprise a more viable strategy to alleviate cognitive impairment than simultaneously targeting individual kinases. Here, we show that treatment with ISRIB, a small molecule compound that targets ISR downstream of eIF2α, restores synaptic plasticity and long-term memory defects in AD mouse models.

ISRIB has been shown to improve symptoms in rodent models of traumatic brain injury, vanishing white matter disease, Down syndrome, SOD1-linked amyotrophic lateral sclerosis (*38*), and prion disease (*18, 19, 22, 23, 39*). However, two previous studies have reported no beneficial effects of ISRIB in AD model mice (*40*). We note that, in contrast to our approach, the previous studies treated AD model mice with a single high-dose i.p. injection of ISRIB (2.5-5 mg/kg). Briggs *et al*. (2017) further reported that prolonged treatment of Tg2576 mice with 5 mg/kg ISRIB led to increased mortality (*40*), which differs from our approach to treat mice with a low-dose of ISRIB (0.25 mg/kg) for several days, which resulted in its detection in the plasma and brains of mice (as shown by LC-MS/MS). Using this dosing regimen, we demonstrate beneficial effects of ISRIB on synapse function and cognitive performance in two different mouse models of AD.

ISRIB was originally described as a memory-enhancing compound (*27*), and its mechanism of action has now been extensively investigated. ISRIB binds to a regulatory site and stabilizes eIF2B, stimulating its GEF activity independently of eIF2α-P (*29*), thus favoring translational derepression. To the best of our knowledge, no off-target effect of ISRIB has been reported.

Our ISRIB dosing regimen (low dose for several days) effectively restored eIF2α-dependent translational control in AD mice, without noticeable biochemical or cognitive effects in WT control mice. Differences between our results and those of Sidrauski *et al*. (*27*) may be due to the fact that, in their case, a single and higher dose of ISRIB (5 mg/kg) was administered within a critical window for memory consolidation, during which de novo protein synthesis is essential for the persistence of newly formed memories (*41*). This is in line with previous reports showing that ISRIB modifies the ISR under conditions of increased eIF2α-P, but not under basal conditions (*42*), (*43*).

We found that ISRIB had a mild effect on aberrant hippocampal *Atf4* mRNA expression in AβO-infused mice. Although not a canonical ISR event, increased *Atf4* mRNA expression has been described as a consequence of ISR under some circumstances (*44*) (*45*). Altogether, these data are consistent with the notion that ISRIB acts downstream to eIF2α-P, restoring AD-linked molecular/cellular defects that are likely driven by chronic low-grade activation of ISR.

Our findings indicate that ISRIB subtly alters amyloid pathology (reduced mean plaque size; increased plaque density) without altering total Aβ_42_ load in the brains of APPswe/PS1ΔE9 mice. One possibility is that the change in amyloid deposits may result from overall resetting of proteostasis induced by ISRIB. Future studies aiming to identify modifications induced by ISRIB treatment in the pool of mRNAs being translated may reveal specific mechanisms of translational control of amyloid deposition.

We found no evidence of decreased microglial activation or astrocyte reactivity in ISRIB-treated APPswe/PS1ΔE9 mice. Thus, it is conceivable that correction of eIF2-dependent translational defects restores memory independently of key hallmarks of the disease (total Aβ accumulation, glial reactivity) raising the prospect that boosting this mechanism may improve cognitive function in patients already exhibiting noticeable neuropathology.

ISRIB rescued impaired synaptic plasticity in the hippocampus of APPSwe/PS1ΔE9 mice, in line with data showing that genetic correction of eIF2α-P levels in neurons restores synaptic plasticity in APPswe/PS1ΔE9 mice (*16*). Moreover, we found that ISRIB rescued reduced dendritic spine densities in both AβO-infused mice and APPSwe/PS1ΔE9 mice. These observations suggest that the beneficial effects of ISRIB-induced translational derepression on memory are linked to improved synapse stability and function.

Our results further demonstrate a selective reduction in eIF2B subunits in AD brains, consistent with a recent single-nucleus transcriptomic study showing that *EIF2B5* expression is reduced in AD neurons (*46*). Although we cannot yet ascertain to what extent the reduction in eIF2B subunits affects protein synthesis in AD, it is reasonable to expect that GDP recycling and translation are impaired under those conditions. Thus, ISRIB might still be effective by stimulating eIF2B assembly (*47*) and the GEF activity of the remaining pool of eIF2B. This is consistent with our observation that acute treatment with ISRIB rescued LTP in hippocampal slices from APPswe/PS1ΔE9 mice.

In conclusion, our findings establish eIF2α-P-dependent translational repression as a potentially druggable target in AD, and open new avenues to investigate the molecular mechanisms by which dysfunctional eIF2 signaling contributes to translational repression in AD. Because the solubility of ISRIB might pose limits to its clinical application, strategies aimed to attenuate the ISR or increase eIF2B activity by using novel eIF2B activators (*22*) or repurposed compounds (*48*) may prove effective to target brain protein synthesis and ward off cognitive decline in AD.

## Materials and Methods

### Ethics

All experiments with mice were carried out in accordance with the “Principals of Laboratory Animal Care” (US National Institute of Health) guidelines and performed under protocols approved and supervised by the Institutional Animal Care and Use Committees of the Federal University of Rio de Janeiro (protocol number IBQM039/16) and New York University (protocol number 09-1332).

### Postmortem human brain tissue

Brain samples (prefrontal cortex) from non-cognitively impaired and AD patients (defined by neuropathological criteria) were obtained from the Emory University Brain Bank. Experimental procedures involving human tissue were in compliance with the NYU Institutional Review Board (IRB). Specimen information can be found in Supplementary Table 1. Post-mortem interval (PMI) varied among donors, but average PMI was similar in AD versus control brains.

### Animals

For the i.c.v. AβO-infusion model, we used three-month-old male and female C57BL/6 mice, equally divided amongst experimental groups. To assess any potential sex bias, we analyzed all groups divided by sex. Mice were obtained from the animal facility at the Federal University of Rio de Janeiro and were maintained on a 12 h light/dark cycle with food and water *ad libitum*, with 5 mice per cage. Transgenic APPswePS1ΔE9 mice (B6/C3 background, JAX#34829) and WT littermates were obtained from the New York University animal facility. Male and female mice (10-13 month-old) were used, and results were analyzed grouping both sexes together. Mice (4 to 5 per cage) were kept on a 12 h light/dark cycle with food and water *ad libitum*. For every experiment, mice were pseudo-randomized into the different experimental groups, and allocation of animals of the same cage in one experimental condition was avoided.

### Reagents

ISRIB (> 98% purity) was from Sigma-Aldrich. Aβ_1-42_ was purchased from California Peptide. Culture media and IR dye-conjugated secondary antibodies were from Li-Cor. Streptavidin-HRP, Streptavidin-AF594, ProLong anti-fade reagent, Tris-glycine gels, and Aβ_42_ ELISA kits were from Invitrogen. Anti-puromycin antibody (12D10) was from EMD Millipore. Anti-ATF4, anti-GFAP, anti-eIF2Bα, anti-eIF2Bε, anti-eIF2γ, anti-β-actin antibodies, and BDNF ELISA kits were from Abcam. Anti-eIF2α-P(Ser51), anti-eIF2α and anti-β-tubulin were from Cell Signaling Technology. 6E10 antibody was from Enzo Life Sciences. Biotin-conjugated anti-Iba1 was from Wako Fujifilm. BCA protein assay kit was from Thermo Fisher. Laemmli Buffer (4x) was from Bio-Rad. Alkyne-conjugated biotin and Protein Reaction Buffer kit were from Click Chemistry Tools.

### ISRIB determination by LC-MS/MS

Analysis of plasma and brain samples were conducted on a QTRAP 4500 triple quadrupole mass spectrometer (Applied Biosystems SCIEX) in the positive ion mode and interfaced with an Ekspert ultraLC 100-XL UHPLC System (Eksigent). Calibration standards (0.003 to 10 μM ISRIB) and quality controls (0.02, 0.2 and 2.0 μM) were prepared in naïve mouse plasma in parallel with mouse plasma study samples (60 μL) by precipitation with three volumes of ice-cold internal standard solution (acetonitrile containing 20 μM theophylline). Precipitated samples were centrifuged at 6,100 x g for 30 min at 4 °C. Following centrifugation, an aliquot of each supernatant was transferred to an autosampler vial and diluted with two volumes of aqueous mobile phase (0.2% formic acid in water). Samples were injected onto a reverse phase analytical column (YMC Triart C18; 2.0 × 50 mm; 1.9 μm; YMC CO) and eluted with a gradient of 0.2% formic acid in acetonitrile. ISRIB and internal standard were monitored by a multiple reaction monitoring (MRM) experiment using Analyst software (v1.6.2, Applied Biosystems SCIEX). Quantitation was conducted using MultiQuant software (v2.1, Applied Biosystems SCIEX) and the resulting calibration curve was fitted by linear regression and 1/x weighting. The lower limit of quantitation (LLOQ) was 0.003 μM.

### Preparation and characterization of amyloid-β oligomers (AβOs)

AβOs were prepared as described (*49*), and each and every preparation was characterized by size-exclusion chromatography (SEC-HPLC), as previously described (*15, 31, 32, 50*). AβO preparations were kept at 4 °C and used within 48 h of preparation.

### Primary neuronal cultures

Pregnant mice were euthanized on the 19^th^ day of gestation, and embryonic brains were collected. Hippocampi and cortices were dissected and homogenized in HBSS containing 0.37% D-glucose. Tissue lumps were removed with a cell restrainer, the cell suspension was centrifuged at 1,500 rpm/5 min and the pellet was re-suspended in DMEM containing 10% fetal bovine serum (FBS). Cell number was estimated using a Neubauer chamber. Cells were plated in (poly-L-lysine + laminin)-coated plates and incubated at 37 °C in a 5% CO_2_ atmosphere for 1h. DMEM was then replaced by Neurobasal medium supplemented with Pen/Strep, Glutamax and 2% B27. Cultured primary neurons were maintained at 37 °C in a 5% CO_2_ atmosphere for 10 days for maturation before experiments. Cells were treated on DIV 12-13 with 0.5 μM AβOs (or vehicle) in the absence of presence of 0.2 μM ISRIB, prepared as previously described (*19*).

### Western blotting

Human and mouse brain tissue samples were homogenized by sonication in RIPA buffer + Halt Protease/phosphatase inhibitor cocktail. Homogenates were centrifuged at 10,000 x *g*/10 min at 4 °C, and the supernatant was harvested for Western blot analysis. Protein concentration was determined by BCA, following manufacturer’s instructions. Protein samples were prepared to a final concentration of 5 μg/μl in 4x Laemmli buffer (Bio-Rad) containing 25 mM DTT. Samples were boiled for 3 min and 50 μg total protein were loaded per lane and resolved in 4-12% SDS-containing Tris-glycine gels. Proteins then were transferred to PVDF or nitrocellulose membranes. Primary antibodies were used in the following dilutions: 1:10,000 anti-β-actin; 1:2,500 anti-puromycin; 1:1,000 anti-ATF4, anti-eIF2α, anti-eIF2α-P; 1:500 anti-eIF2Bε, anti-eIF2Bα, anti-eIF2γ. Secondary antibodies, either IR dye-conjugated (Li-Cor) or horseradish peroxidase (HRP)-conjugated, were used at 1:10,000 dilution. ECL plus was used to develop HRP-labeled blots.

### RT-PCR

Total RNA was extracted from cultures using the SV Total RNA Isolation System (Promega), following manufacturer’s instructions. RNA concentration and purity were determined by absorption at 260 nm and 280 nm. For quantitative real-time reverse transcription PCR (qRT–PCR), 1 μg of total RNA was used for complementary DNA (cDNA) synthesis using the High Capacity cDNA Reverse Transcription kit (ThermoFisher Scientific). Quantitative expression analysis of targets was performed on a 7500 Fast Real-Time PCR system (ThermoFisher Scientific) with the Power SYBR Green PCR Master Mix. β-Actin (actb) was used as an endogenous reference gene for data normalization. qRT–PCR was performed in 15 μl reaction volumes. Primer sequences used for Atf4 and Actb amplification were: Fwd Atf4 – CCACCATGGCGTATTAGAGG; Rev ATF4 – CTGGATTCGAGGAATGTGCT; Fwd Actb – TGTGACGTTGACATCCGTAAA; and Rev Actb – GTACTTGCGCTCAGGAGGAG. Cycle threshold (CT) values were used to calculate fold-changes in gene expression using the 2^−ΔΔCt^ method (*51*).

### ELISA

Tissue samples were homogenized in 100 mM Tris, 150 mM NaCl, 1 mM EGTA, 1 mM EDTA, and 1% Triton X-100, supplemented with protease and phosphatase inhibitor cocktail (ThermoFisher). ELISAs for BDNF and Aβ_42_ were run following manufacturer’s instructions.

### BONCAT

Bio-Orthogonal Noncanonical Amino Acid Tagging (BONCAT) was utilized for de novo protein synthesis assessment (*52*). Primary neuronal cultures had their medium replaced by methionine-free RPMI supplemented with 2% B27, 1% Glutamax and 0.25% glucose. Cells were incubated for 30 min to allow for methionine starvation and azidohomoalanine (AHA), a methionine analog azide, was added at 1 mM final concentration to label newly synthesized proteins. AβOs (0.5 μM) and/or ISRIB (0.2 μM) were added, and cells were incubated for 2 h at 37 °C. Cellular content was harvested using RIPA buffer (Pierce) supplemented with Halt Protease/Phosphatase inhibitor cocktail. Protein levels were determined using the BCA method (Thermo Pierce). AHA-conjugated proteins were biotinylated using the Protein Reaction Buffer kit (Thermo Fisher), following manufacturer’s instructions. Newly synthesized proteins were precipitated using streptavidin-conjugated resin and were detected by Western blotting using streptavidin-conjugated HRP (1:2,000).

### SUnSET

Hippocampal slices were prepared as previously described (*16*). Briefly, 400 μm hippocampal slices were obtained using a chopper, and recovered in artificial cerebrospinal fluid (aCSF) for 1h. Slices were then exposed to 1 μM AβOs (or vehicle) in the absence or presence of 0.2 μM ISRIB for 3h. Puromycin was added to the media to a final concentration of 5 μg/ml during the last hour of the experiment. After incubation, slices were flash frozen and processed for Western blotting. Puromycin incorporation was quantitated as a measure of newly synthesized proteins. The representative image shown in Figure 3J had its brightness level linearly increased solely for visualization purposes, but quantification of incorporated puromycin was performed in raw images without any manipulation.

### Animal treatments

Intracerebroventricular (i.c.v.) infusions of AβOs were performed as described previously (*15, 31, 32, 53*). Mice were anesthetized briefly with 2% isoflurane and AβOs were injected 1 mm to the right of the midline point equidistant of each eye and 1 mm posterior to a line drawn through the anterior base of the eyes. Mice were then placed back in their cages.

Mice were injected daily intraperitoneally with 0.25 mg/kg ISRIB or vehicle (200 μL per injection). Treatment regimen is described in each section of “Results” and followed a protocol previously established in literature, known to have a beneficial effect in a murine model of prion disease (*18*). Using this protocol, Halliday and co-workers (2015) showed that substantial amounts of ISRIB reached the brain and persisted for at least 24h. For salubrinal experiments, mice were concomitantly injected daily with ISRIB (0.25 mg/kg) and salubrinal (1 mg/kg). Results obtained in this study were used as a template for subsequent studies using AD mouse models.

### Behavioral analyses

Long-term memory was assessed using the Novel Object Recognition (NOR), Contextual Fear Conditioning (CFC), and Morris water maze (MWM) tests, as described below. Locomotor activity was assessed in an Open Field Arena (OFA). In all behavioral experiments, the experimenter was blinded to the groups tested. Animals that had biased exploration during training in the NOR (100% exploration of a specific object during training phase) were excluded from analysis. In the MWM, mice that did not swim during trials were excluded from both the learning analysis and the probe trial. Finally, in the CFC task, mice with unaltered behavior after being shocked were excluded from the analysis.

### Novel Object Recognition

Tests were carried out using a NOR box (30 × 30 × 50 cm). In the training phase, mice were exposed to two identical objects, which they were allowed to freely explore for 5 minutes. Time spent exploring each object was recorded and twenty-four hours later, in the test phase, one of the objects was replaced by a novel object. Mice again were exposed to the objects for 5 minutes, and the total exploration time of both old (familiar) and new (novel) objects were determined. Discrimination index was determined by: (T_novel_ − T_familiar_) / (T_novel_ + T_familiar_). After the task was completed, mice were placed in a different arena for 5 minutes, where total distance, average velocity and total time spent at the periphery were recorded and determined using ANY-Maze software (Stoelting Co.).

### Contextual Fear Conditioning

To assess contextual fear memory, a two-phase protocol was used, as described (*15, 54*) with minor modifications. In the training phase, mice were presented to the conditioning cage (40 × 25 × 30 cm), which they were allowed to freely explore for 2 minutes followed by application of a single foot shock (0.35 mA) for 2 s. Mice were kept for another 30 s in the cage and removed. On the next day, mice were presented to the same cage for 5 minutes without receiving a foot shock. Freezing behavior was recorded automatically using the Freezing software (Panlab) and was used as a memory index. For APPSwe/PS1ΔE9 mice, due to the extensive memory impairment in these mice, protocol included 2 min of free exploration, followed by two foot shocks (0.8 mA), 2 s each, spaced by 30 s. Mice were then kept for another 2 min in the cage and removed.

### Morris water maze

The Water Maze test was carried out as previously described (*16, 55*). Briefly, 3 training sessions per day were carried out for 5 days, except in the first day, when mice were trained in 4 sessions. Time to reach the platform was recorded as a measure of learning capacity. Average velocity and total distance traveled in the pool were recorded as control locomotor parameters. Probe trials were performed for 1 minute 24 h after the last training session. Time spent in the target quadrant was recorded as a memory output. Recording was performed using Ethovision software.

### Open Field Arena

Locomotor activity in APPswePS1ΔE9 mice was assessed using the OFA (Med Associates Inc). Mice were allowed to freely explore the box for 15 min, during which their movement was recorded using Activity MDB software (Med Associates Inc). Total distance traveled, average velocity and total time spent at the periphery were recorded.

### Golgi staining

Mice were euthanized and whole brains were carefully harvested, rinsed in PBS and prepared for Golgi staining using the FD Rapid Golgi Kit (FD Neurotechnologies) following manufacturer’s instructions. Briefly, brains were immersed in a mixture containing solutions A and B (1:1) for 14 days protected from light, with gentle swirling twice a week. Brains were then incubated in solution C for 72 hours before sectioning. After staining, brains were sliced on a Leica Vibratome in 200 μm-thick sections and developed for 5 minutes in solutions D and E, followed by dehydration. Slices were mounted on gelatin-coated slides and imaged on a Nikon Eclipse TE2000-U microscope under bright field illumination. For dendritic spine quantification, 3 dendrite segments of 10-20 μm that were at least 50 μm from the cell soma were selected per neuron. Five neurons were analyzed per brain. Total dendritic spines in each segment were counted and normalized by total length of the dendritic segment. For image analysis, the experimenter was blinded to the conditions analyzed. For visualization purposes, representative images shown in Figure 3E had their brightness and contrast linearly altered, but spine counting was performed in the absence of any image manipulation.

### Electrophysiology

Acute 400 μm transverse hippocampal slices from WT or APPswe/PS1ΔE9 mice were prepared using a Leica VT1200S vibratome as described previously (*56*). Slices were maintained at room temperature for at least 2 h in artificial cerebrospinal fluid (aCSF) containing (in mM) 118 NaCl, 3.5 KCl, 2.5 CaCl_2_, 1.3 MgSO_4_, 1.25 NaH_2_PO_4_, 5.0 NaHCO_3_ and 15 glucose, bubbled with 95% O_2_ / 5% CO_2_. For electrophysiology, monophasic, constant-current stimuli (100 μs) were delivered with a bipolar silver electrode placed in the stratum radiatum of area CA3. Field excitatory postsynaptic potentials (fEPSPs) were recorded using a glass microelectrode from the stratum radiatum of area CA1. Late-LTP (L-LTP) was induced using high-frequency stimulation (HFS) consisting of two 1-sec 100 Hz trains separated by 60 sec, each delivered at 70-80% of the intensity that evoked spiked fEPSPs. Slices were perfused with ISRIB (0.2 μM in ACSF) or vehicle during the whole recording process.

### Immunohistochemistry

WT and APPSwe/PS1ΔE9 treated with ISRIB (as described above) were anesthetized with a ketamine/xylazine mixture. Mice were first perfused with PBS for 45 seconds, and then with 4% PFA for 90 seconds. Brains were carefully removed from the skull and post-fixed in 4% PFA for 48h at 4 °C, under agitation. Brains were blocked on 3% agarose and sectioned using a VT1200 S vibratome (Leica Biosystems), to 40 μm sections. Sections were kept at 4 °C until use. For Aβ (6E10), Iba1 and GFAP staining, free-floating sections were initially permeabilized using 0.5% Triton X-100 solution for 15 min. Sections were then blocked for 1h with 5% NGS, and incubated overnight with 6E10 (mouse, 1:200) and anti-GFAP (chicken, 1:1000) solution. Sections were washed 3 times using 0.1% Triton X-100 and incubated with anti-rabbit and anti-chicken secondary antibodies (1:500) for 90 min. Sections were again washed 3 times with 0.1% Triton X-100 and subsequently incubated overnight with biotin-conjugated anti-Iba1 (rabbit, 1:500). Sections were washed 3 times using 0.1% Triton X-100 and incubated with Alexa Fluor 594-conjugated streptavidin (1:500) for 90 min. Sections were washed 3 times with 0.1% Triton X-100 and mounted using Prolong with DAPI. Imaging was performed on a Leica SP8 confocal microscope. Images were acquired and analyzed using identical parameters across the experiment. Total fluorescence was determined by generating a mask in Fiji. The mask was used to determine the threshold separating signal and noise. Total pixel intensity (determined by the “Raw Integrated Density” variable) was then obtained using the whole image as region of interest (ROI). For glial content surrounding plaques, 20X magnification images were used. A region containing the plaque and an ~30 μm radius surrounding area was used as ROI. Total Iba1 or GFAP fluorescence were determined as above. For image analysis, the experimenter was blinded to the conditions.

### Statistical analysis

Pilot experiments were performed to allow calculations of sample and effect sizes by power analysis. G Power software (Düsseldorf University) was used for power analysis calculations after determination of the statistical analysis that would be used to analyze each experiment. Statistical analyses were performed using GraphPad Prism 6 software. Differences between two independent groups were analyzed using Student’s t-test. When three or more independent experimental groups were compared, ANOVA was used, followed by appropriate *post-hoc* tests, as stated in “Figure Legends” and in Supplemental Table 2.

## Supplementary Materials

**Figure S1.** ISRIB does not attenuate the increase in hippocampal Atf4 mRNA, or the reductions in BDNF, PSD-95 and synaptophysin induced by AβOs in mice.

**Figure S2.** ISRIB does not alter locomotor activities.

**Figure S3.** ISRIB restores GADD34 levels in the hippocampus of APPSwe/PS1ΔE9 mice.

**Figure S4.** Locomotor/exploratory activities and body weight of APPswe/PS1ΔE9 are not altered by ISRIB treatment.

**Figure S5.** Gliosis in APPSwe/PS1ΔE9 hippocampus (dentate gyrus/stratum lacunosum moleculare) is unaffected by ISRIB treatment.

**Fig S6.** Full Western blots used in the current study.

**Supplementary Table 1.** Demographics of control and AD brains.

## Acknowledgements

We thank Dr. Anna Vorobyeva (NYU) for insightful discussions, Dr. Maggie Mamcarz (NYU) for APPSwe/PS1ΔE9 genotyping, and Dr. Ronir Raggio Luiz (Federal University of Rio de Janeiro) for assistance with statistical analyses.

## Author contributions

M.M.O., M.V.L., E.K. and S.T.F. designed the study. M.M.O., M.V.L., F.L, N.K., W.Y., G.U., D.D.P.F. and P.H.J.M. performed research. M.M.O., M.V.L., F.L, N.K., W.Y., G.U., D.D.P.F. and P.H.J.M. analyzed data. T.M., S.B., F.G.D.F, E.K. and S.T.F. contributed reagents, materials, animals and analysis tools. M.M.O., M.V.L., T.M., E.K., and S.T.F. analyzed and discussed results. M.M.O., E.K., and S.T.F. wrote the manuscript with input from other authors.

---

**Figure S1.**
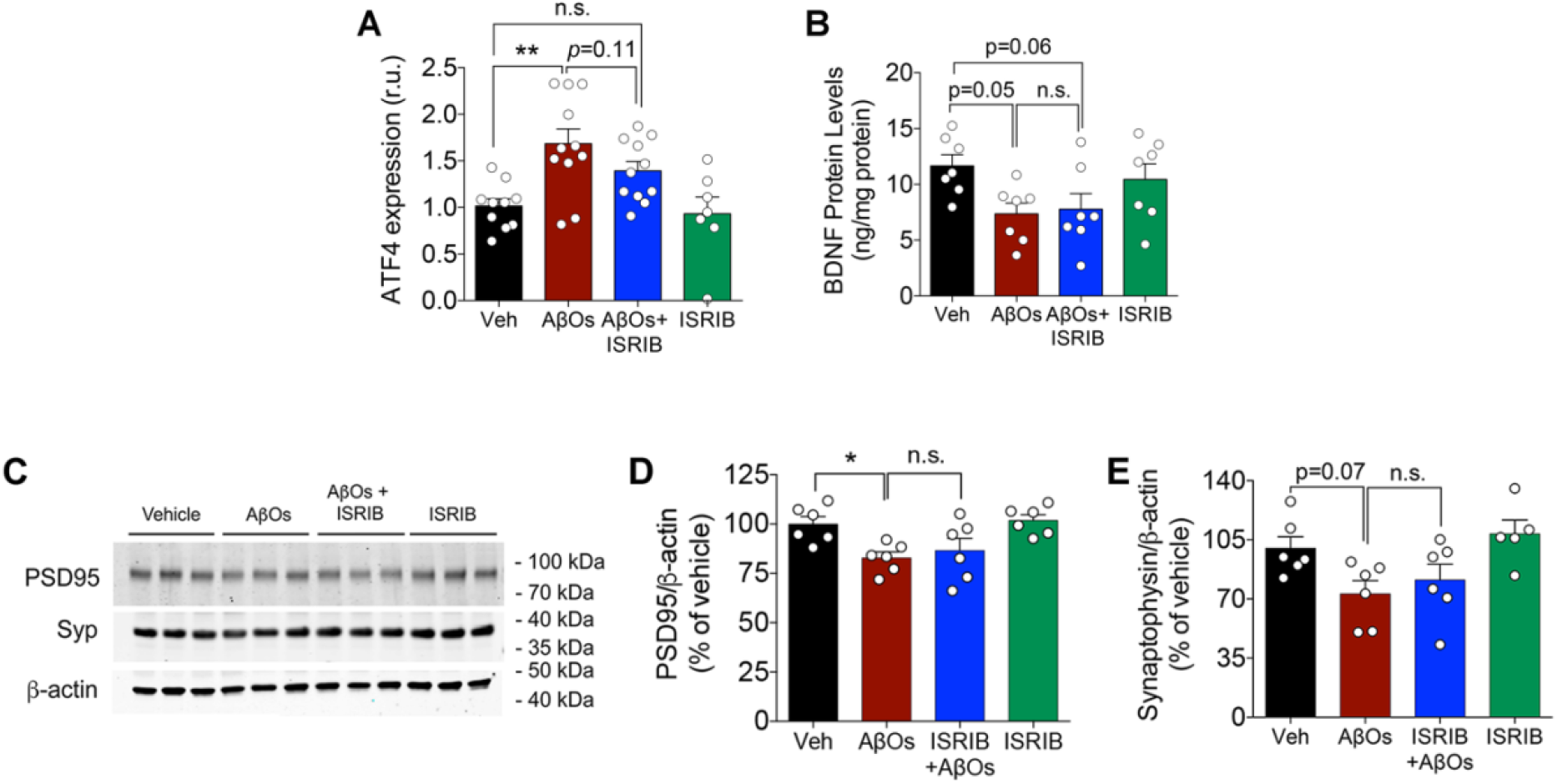
ISRIB does not attenuate the increase in hippocampal ATF4 mRNA, or reductions in BDNF, PSD-95 and synaptophysin protein levels induced by AβOs in mice. Mice received an i.c.v. infusion of 10 pmol AβOs (or vehicle) and were treated daily with ISRIB (0.25 mg/kg, i.p., for 5 days). Hippocampal tissue was collected 12 days after AβO infusion. (A) ATF4 mRNA levels measured by qPCR in the hippocampi of AβO-infused mice treated or not with ISRIB (N = 7-11 mice/group). (B) BDNF measured by ELISA in the hippocampi of the same mice (N = 7 mice/group). (C-E) PSD-95 and synaptophysin levels evaluated by Western blotting in hippocampi of AβO-infused mice and normalized by β-actin levels (N = 6 mice/group). Veh = vehicle. Two-Way ANOVA followed by Dunnet’s *post-hoc* test, \*\**p* < 0.01. Dots represent individual mice.

**Figure S2.**
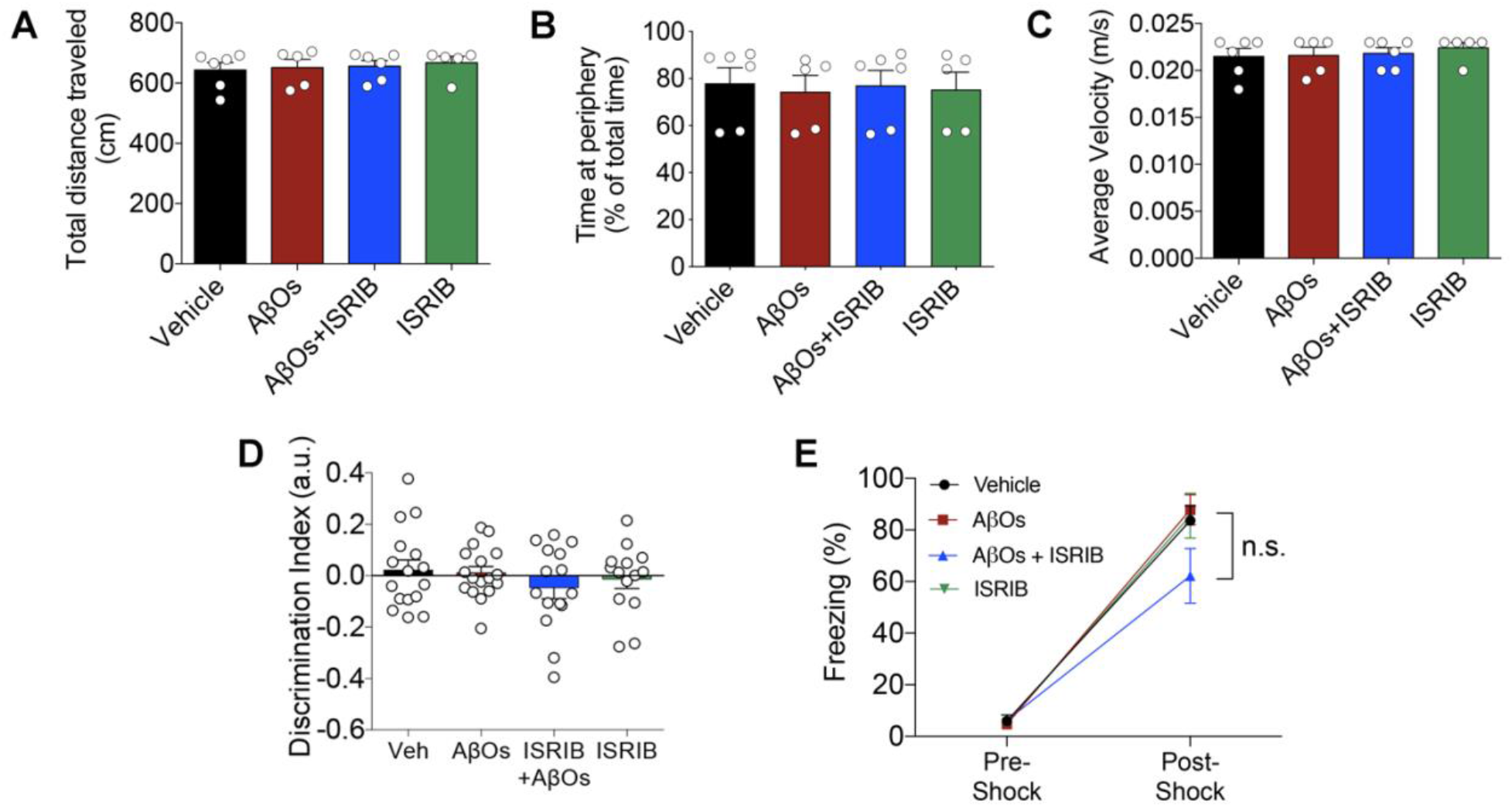
ISRIB does not alter locomotor activities. Mice received an i.c.v. infusion of 10 pmol AβOs and/or were daily treated with ISRIB (0.25 mg/kg; i.p.) as described in Figure 3A. They were allowed to freely explore an open field arena for 5 min, during which total distance traveled (A), time spent at the periphery (B) and average velocity (C) were recorded (N = 5-6 mice/group). (D) Discrimination index of the NOR training session (N = 14-17 mice/group). (E) Freezing behavior during the FC training session, divided in periods pre-shock and post-shock (N = 6-8 mice/group). Veh = vehicle. Two-Way ANOVA followed by Dunnet’s *post-hoc* test. Dots correspond to individual mice.

**Figure S3.**
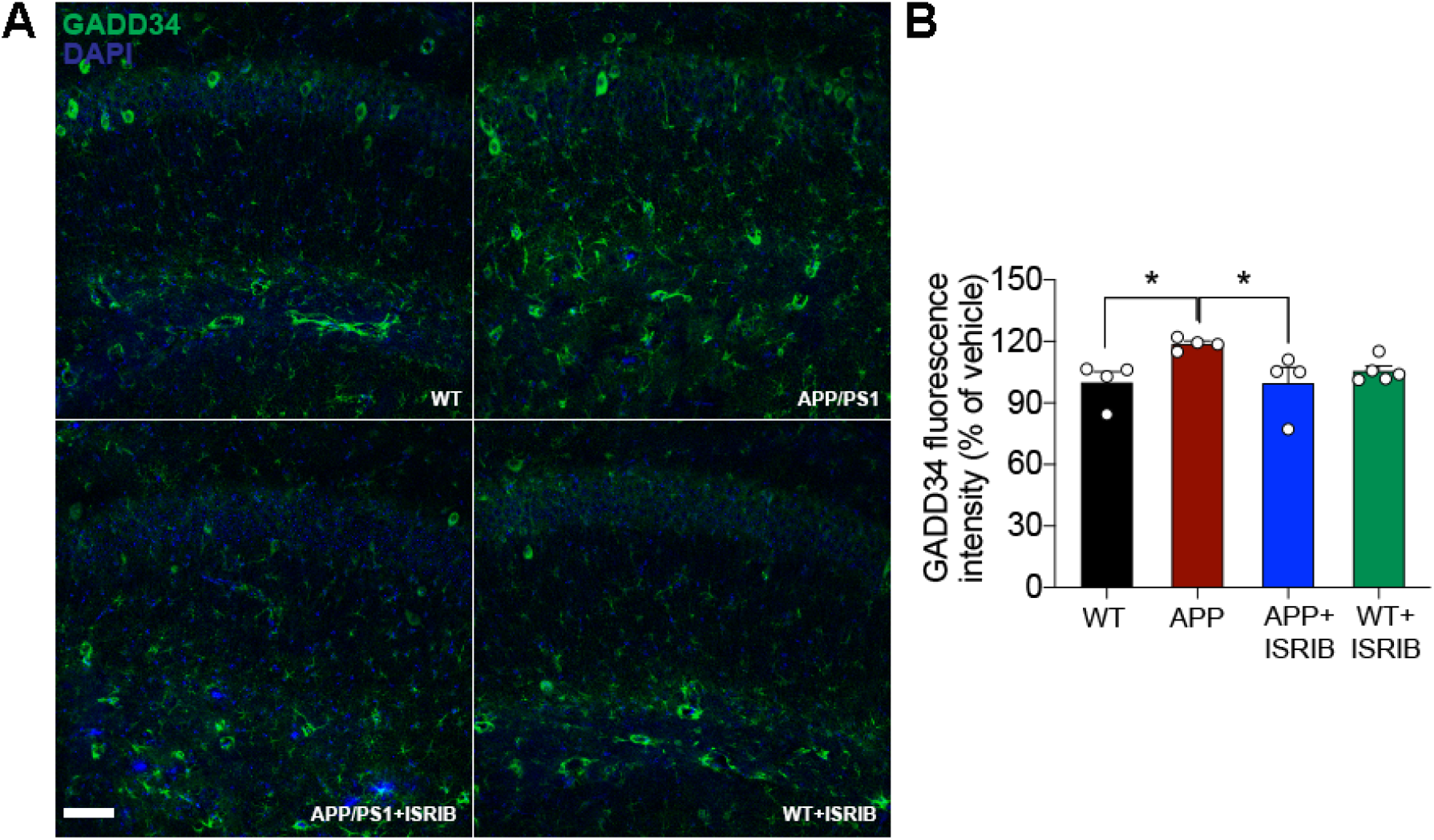
ISRIB rescues GADD34 levels in the hippocampus of APPSwe/PS1ΔE9 mice. (A) Representative images of GADD34 levels (green) in the CA1 region from the hippocampus of WT or APPSwe/PS1ΔE9 mice, treated or not with ISRIB (0.25 mg/kg). Scale bar = 20 μm. (B) Quantification of total levels of GADD34 levels in the CA1 region (N = 4-5 mice/group). * = *p* < 0.05, Two-Way ANOVA followed by Dunnet’s test. Dots represent individual mice.

**Figure S4.**
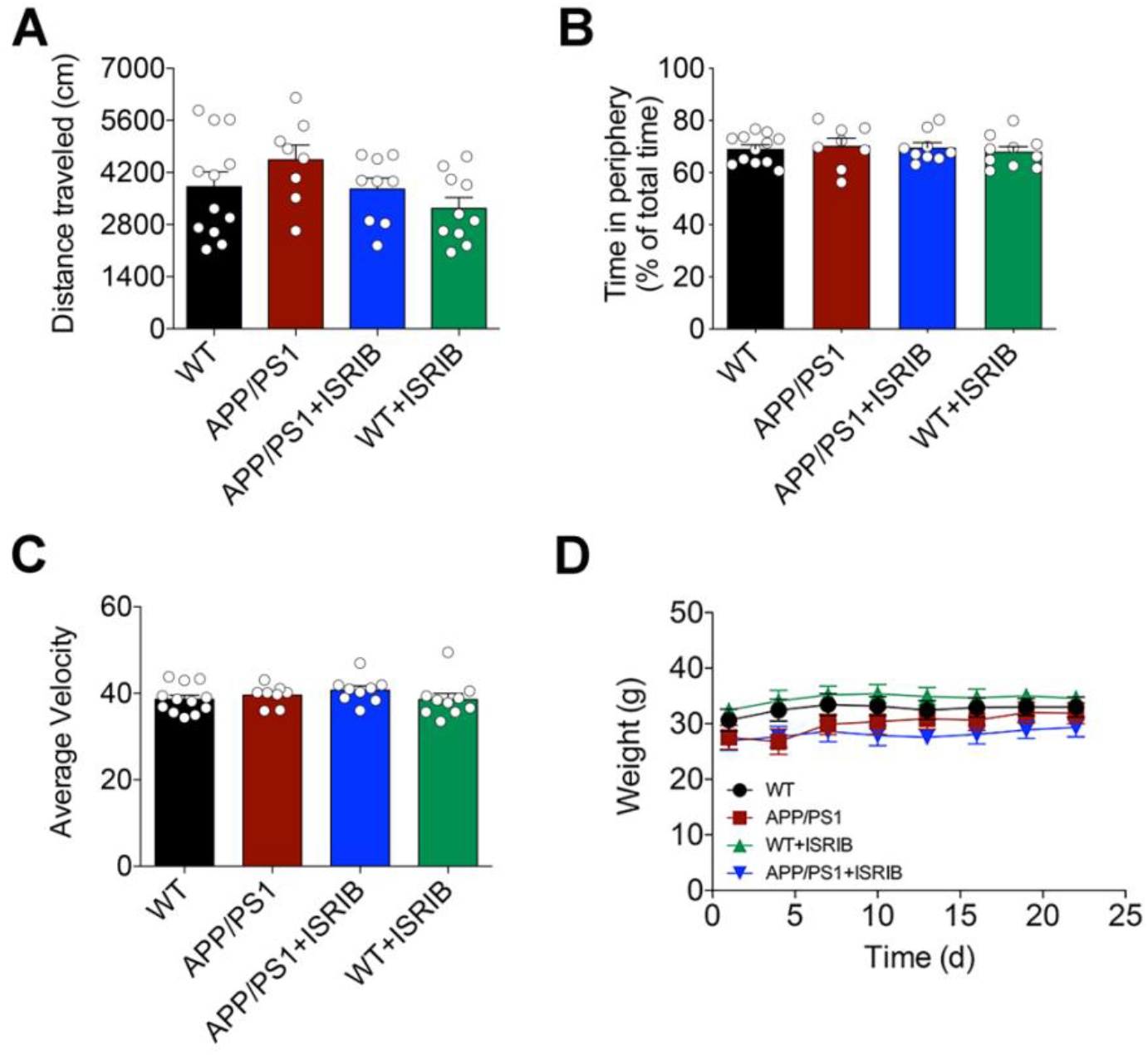
Locomotor/exploratory activities and body weight of APPswePS1ΔE9 are not altered by ISRIB treatment. APPswePS1ΔE9 (or WT littermates) mice were treated with ISRIB (0.25 mg/kg, i.p., daily) for 2 weeks prior to behavioral assessment, and during the battery of behavioral tasks (total time of treatment = 24 days). To assess motor and exploratory behaviors, mice were allowed to freely explore an open field arena for 15 min. Total distance traveled (A), percentage of time at the center of the arena (B) and average velocity (C) were measured (N = 8-11 mice/group). (D) Mice were weighted every 3 days for 21 days of treatment (N = 8-11 mice/group). Dots represent individual mice.

**Figure S5.**
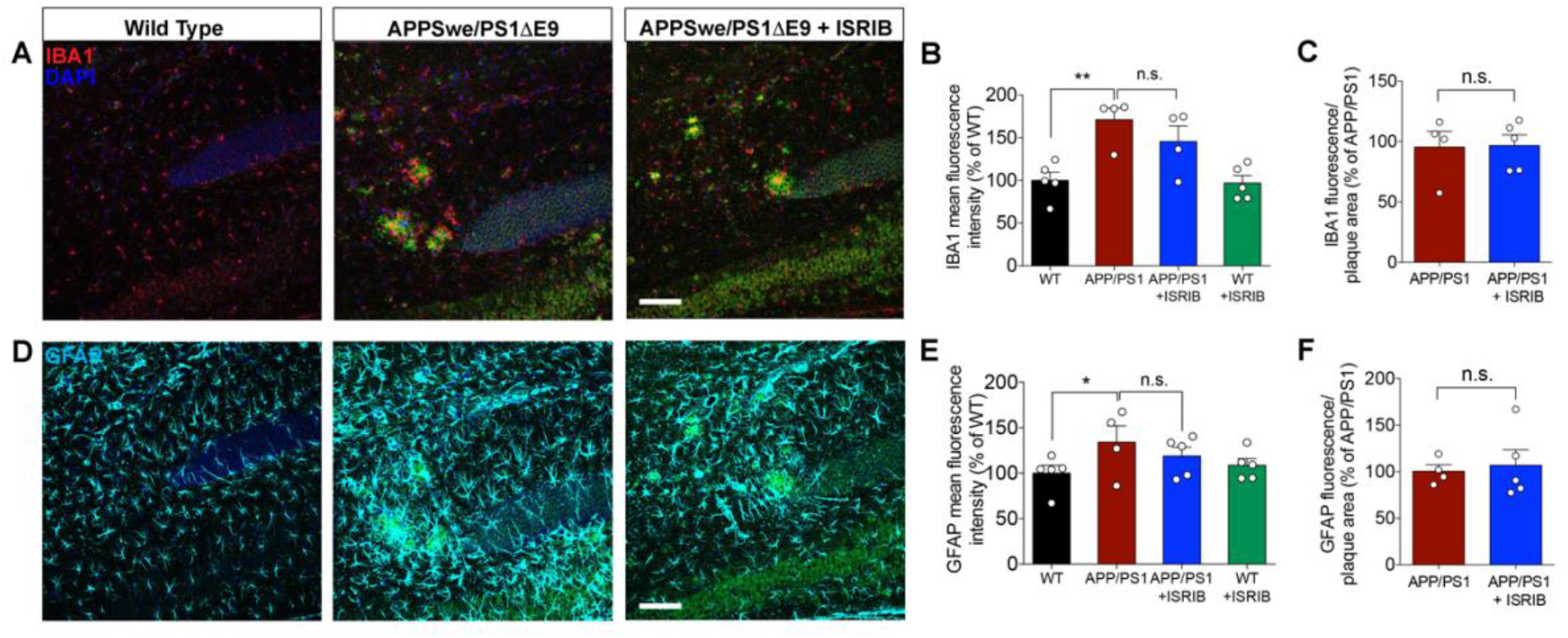
Gliosis in APPSwe/PS1ΔE9 hippocampus (dentate gyrus/stratum lacunosum moleculare) is unaffected by ISRIB treatment. (A) Representative Iba-1 immunoreactivity images (red) in the hippocampi of WT, APPSwe/PS1ΔE9 or ISRIB-treated APPSwe/PS1ΔE9 mice (scale bar = 50 μm). (B) Quantitative analysis of Iba-1 immunoreactivity (mean fluorescence intensity; N = 4-5 mice/group). (C) Quantitative analysis of Iba-1 immunoreactivity (mean fluorescence intensity) co-localized with or immediately surrounding plaques (N = 4-5 mice/group). (D) Representative GFAP immunoreactivity images (cyan) in the hippocampi of W, APPSwe/PS1ΔE9 or ISRIB-treated APPSwe/PS1ΔE9 mice (scale bar = 50 μm). (E) Quantitative analysis of GFAP immunoreactivity (mean fluorescence intensity; N = 4-5 mice/group). (F) Quantitative analysis of GFAP immunoreactivity (mean fluorescence intensity) co-localized with or immediately surrounding plaques (N = 4-5 mice/group). n.s. = not statistically significant. *p<0.05, **p<0.01, Two-Way ANOVA followed by Fisher’s test. Dots represent individual mice.

**Fig S6.**
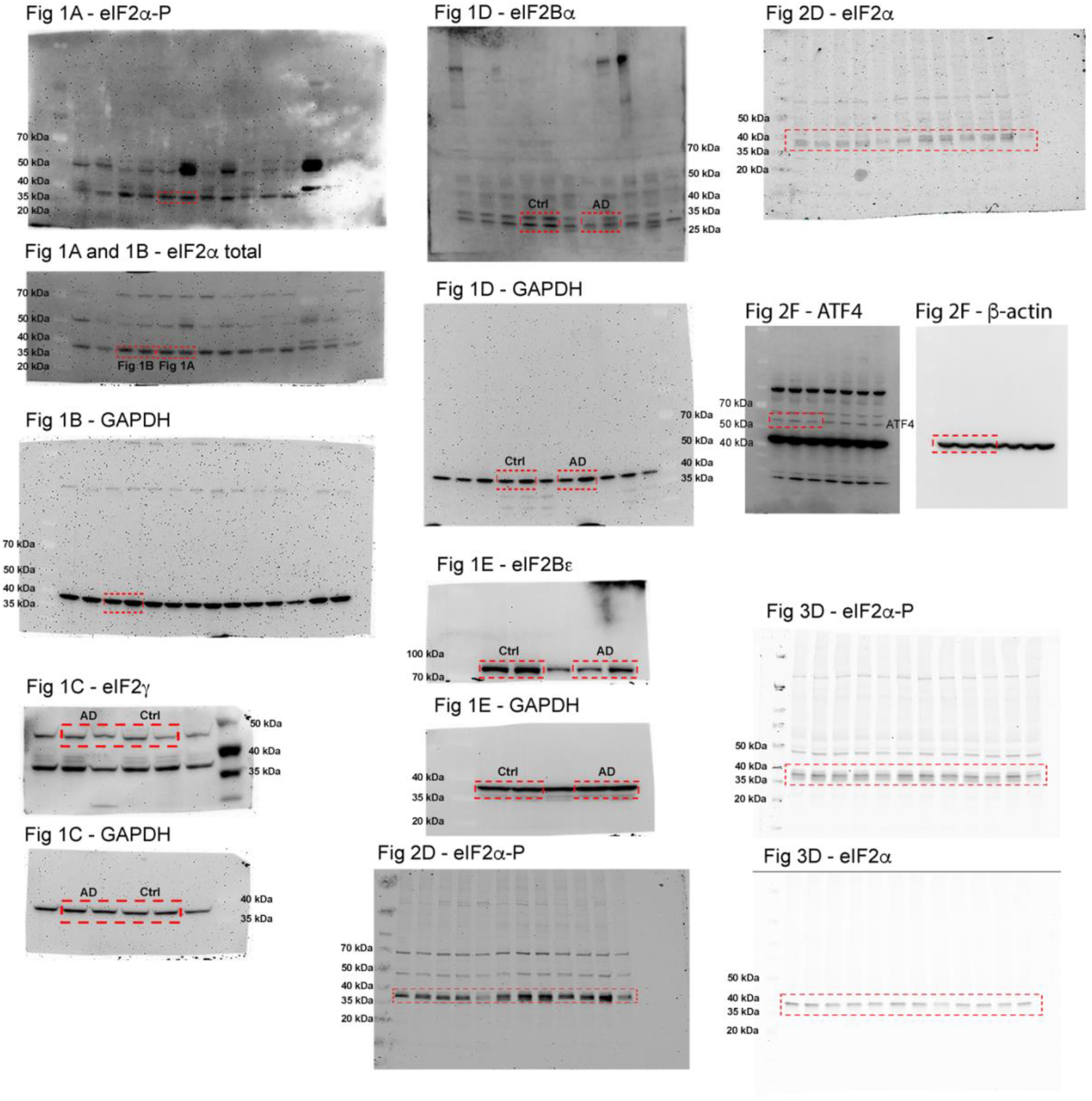
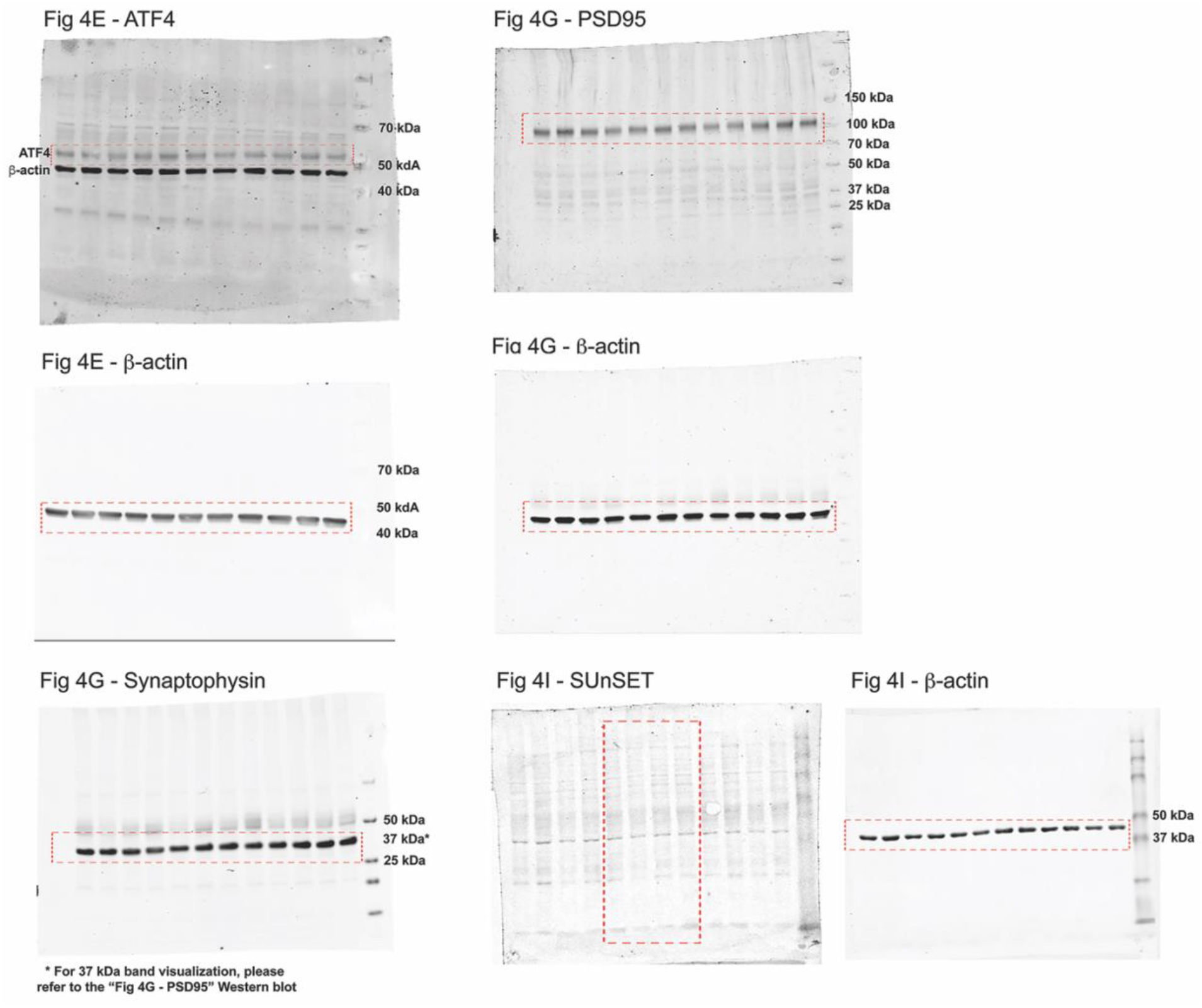
Full Western blots used in the current study. Lanes shown in main figures are boxed in red.

**Supplementary Table 1.** Demographics of control and AD brains.

| Case Number | Primary Neuropathologic Diagnosis | PMI (hr) | Age at Onset | Age at Death | ApoE | Sex |
| --- | --- | --- | --- | --- | --- | --- |
| OS02-252 | AD | 17 |  | 50 | E3/3 | Female |
| OS00-11 | AD | 4 | 49 | 55 | E3/3 | Male |
| E06-154 | AD | 5 | 52 | 60 | E4/4 | Male |
| OS-140909 | AD | 6 | 51 | 61 | E3/3 | Female |
| OS02-163 | AD | 11 | 53 | 70 | E3/4 | Male |
| E08-53 | AD | 8 | 70 | 78 | E3/3 | Female |
| E10-149 | AD | 6.5 | 77 | 90 | E2/4 | Male |
| E08-108 | AD | 4 | 76 | 94 | E3/4 | Female |
| E05-130 | Control | 3 | N/A | 52 | E3/4 | Female |
| E14-06 | Control | 12.5 | N/A | 56 |  | Male |
| E05-74 | Control | 6 | N/A | 59 | E2/3 | Male |
| E15-106 | Control | 6 | N/A | 61 |  | Female |
| E16-45 | Control | 2.5 | N/A | 70 | E3/3 | Male |
| E08-101 | Control | 11.5 | N/A | 78 | E3/3 | Female |
| E13-27 | Control | 6 | N/A | 91 | E3/3 | Female |
| E10-142 | Control | 5.5 | N/A | 94 | E3/3 | Male |

PMI: post-mortem interval
ApoE: Apolipoprotein E alleles

